# The axon initial segment drives the neuron’s extracellular action potential

**DOI:** 10.1101/266734

**Authors:** Douglas J. Bakkum, Milos Radivojevic, Marie Engelene J. Obien, David Jaeckel, Urs Frey, Hirokazu Takahashi, Andreas Hierlemann

## Abstract

Extracellular voltage fields produced by a neuron’s action potentials provide a primary means for studying neuron function, yet their biophysical sources remain ambiguous. The neuron’s soma and dendrites are thought to drive the extracellular action potential (EAP), while the axon is usually ignored. However, by recording voltages of single neurons in dissociated rat cortical cultures and Purkinje cells in acute mouse cerebellar slices at hundreds of sites, we find instead that the axon initial segment dominates the EAP, and, surprisingly, the soma shows little or no influence. As expected, this signal has negative polarity (charge entering the cell) and initiates at the distal end. Interestingly, signals with positive polarity (charge exiting the cell) occur near some but not all dendritic branches and occur after a delay. Such basic knowledge about which neuronal compartments contribute to the extracellular voltage field is important for interpreting results from all electrical readout schemes. Moreover, this finding shows that changes in the AIS position and function can be observed in high spatiotemporal detail by means of high-density extracellular electrophysiology.

**Key points summary:**

- The neuron’s soma and dendrites are thought to give rise to its extracellular voltage signal, while signals from the axon are usually considered negligible.
- Instead, we found that the largest amplitude of the extracellular signal originates from the axon initial segment, not from the soma.
- This finding shows that changes in the AIS position and function can be observed in high spatiotemporal detail by means of high-density extracellular electrophysiology.

**Abbreviations:**

ACSF: artificial cerebrospinal fluid
AIS: axon initial segment
AnkG: ankyrin-G
CMOS: complementary-metal-oxide-semiconductor
DMEM: Dulbecco’s modified eagle medium
EAP: extracellular action potential
GAD67-GFP: glutamic acid decarboxylase 67-green fluorescent protein
MAP2: microtubule-associated protein 2.

## Introduction

How different neuronal compartments and different cell types contribute to the extracellular action potential (EAP) remains unclear and without sufficient direct experimental evidence. A variety of theoretical (Rall, 1962; Holt and Koch, 1999; Gold et al., 2006; Pettersen and Einevoll, 2008; Einevoll et al., 2013) and a limited number of experimental (Grace and Bunney, 1983; Henze et al., 2000; Frey et al., 2009; Buzsaki et al., 2012; Baranauskas et al., 2013; Anastassiou et al., 2015; Petersen et al., 2015) studies address the subcellular origins of EAPs. In most, axonal contributions to the EAP have been considered negligible or small, possibly due to a notion that small cell compartments require less current to depolarize (Rall, 1962; Humphrey and Schmidt, 1990). Along these lines, initial patch-clamp data shows similar AIS and somatic sodium currents per area of membrane (Colbert and Johnston, 1996), which are the currents responsible for action potential generation. In other words, the much larger soma would produce a much greater cumulative current. On the other hand, immunohistological evidence shows that sodium channels are expressed to a much high degree in the AIS (Hu et al., 2009). There, they are tightly coupled to the actin cytoskeleton and not easily drawn into a patch pipette, which suggests that the initial patch-clamp data may underestimate sodium current at the AIS (Kole et al., 2008). Recent quantitative freeze-fracture immunogold labeling (Lorincz and Nusser, 2010), outside-out patch clamp recordings (Hu et al., 2009), and experimentally constrained models (Kole et al., 2008) show that the AIS of pyramidal neurons has about 35 to 50 times higher sodium channel density than the soma or proximal dendrites. Likewise, while not focusing on the EAP, early modeling studies predict that a 20- to 500-fold higher density of sodium channels is necessary for action potentials to initiate in the AIS (Dodge and Cooley, 1973; Mainen et al., 1995). A simplistic calculation using a 50-fold increase shows that the AIS would indeed be able to influence the EAP: an AIS would have approximately 5 times greater total sodium conductance than a soma, assuming a 17-μm diameter spherical soma and a 31-μm length cylindrical AIS (mean values from our data, which also matches ratios in (Mainen et al., 1995)) with an estimated 1 μm diameter; conductance is considered to be proportional to surface area multiplied by the channel density. However, whether or not the AIS has more sodium channels than the soma, and to what degree, remains debated (Colbert and Johnston, 1996; Kole et al., 2008; Schmidt-Hieber and Bischofberger, 2010; Hallermann et al., 2012).

Here, we compared neurons’ EAPs to fluorescence images of cell morphology in primary rat cortical cultures and acute mouse cerebellar slice preparations and present direct evidence that the EAP is driven by the AIS, not the soma. The spatial distribution and strength of the EAP is related to the AIS’s geometry and degree of formation. Our results add experimental evidence into the debate about the distribution of sodium channels in a neuron. To gather the data, neurons’ EAPs were recorded in high spatial detail by using custom microelectrode arrays containing 11,011 densely packed electrodes (17.8 μm pitch; 8.2 × 5.8 μm2 area per electrode (Frey et al., 2010)). Immediately after recording, cells were imaged in order to match which cell compartments contributed to the neurons’ EAPs. For the primary cultures, spontaneous EAPs of single neurons were sampled at multiple locations simultaneously at 2 to 3 weeks in vitro; to minimize overlap of voltage signals in recordings and neurites in images between neighboring neurons, we grew low-density cultures that contained a few to tens of neurons per array. A GAD67-GFP knock-in mouse line (Tamamaki et al., 2003) that allowed live imaging of Purkinje cells was used for acute slice recordings. We conclude with a brief discussion about how our results can affect interpretations of extracellular data.

## Methods

### Animal use

Protocols were approved by the Basel Stadt veterinary office according to Swiss federal laws on animal welfare and the Animal Research Committee of the RIKEN Center for Developmental Biology.

### Low-density cell culturing

Cells from embryonic day 18 Wistar rat cortices were dissociated in trypsin, followed by mechanical trituration, and 1000 cells were seeded over an area of ~4 mm2 on top of the microelectrode arrays. Neurons and glia grew in 2 mL of serum-containing Neurobasal-based media for 5 days. Thereafter, half of the media was changed once a week to serum-containing DMEM-based media. Detailed media recipes are in Bakkum et al., 2013.

### Immunocytochemistry

The neural source of extracellular signals was verified optically by immunocytochemistry performed immediately after a recording session. A detailed protocol is given in (Bakkum et al., 2013). The primary antibodies to microtubule-associated protein 2 (MAP2; Abcam ab5392; RRID:AB_2138153) and ankyrin-G (AnkG; NeuroMab clone N106/36; RRID:AB_10697718) were diluted 1:200 and left overnight at 4°C on a shaker set to low speed. The secondary antibodies containing Alexa Fluor 647 (RRID:AB_10374876) or Alexa Fluor 488 (RRID:AB_10561551) were diluted 1:200 and left for 1 hour at room temperature in the dark.

### Acute cerebellar slice preparation

GAD67-GFP knock-in mice (Tamamaki et al., 2003), postnatal day 21-37, were obtained from the Laboratory for Animal Resources and Genetic Engineering in RIKEN Center for Developmental Biology Kobe, Japan. The mice were anaesthetized by isoflurane inhalation and then decapitated. The procedure for preparing cerebellar slices has been adapted from (Frey et al., 2009). Briefly, the brain was dissected and immediately immersed in ice-cold carbogenated artificial cerebrospinal fluid (ACSF) containing in mM: NaCl 125, KCl 2.5, NaH_2_PO_4_ 1.25, MgSO_4_ 1.9, Glucose 20, NaHCO_3_ 25. The cerebellum was separated from the cortex by cutting with a blade and glued onto a tray along its sagittal plane. Parasagittal cerebellar slices (150-200 μm thick) were obtained using a vibratome (Leica VT1200S). The slices were incubated at 35°C in ACSF with 2 mM CaCl_2_ for 30-45 minutes and then maintained at room temperature until recording.

### Recording EAPs with CMOS-based microelectrode arrays

Cortical networks were grown for two to three weeks over 11,011-electrode complementary-metal-oxide-semiconductor-based (CMOS) microelectrode arrays (1.8 × 2.0 mm2 area containing 8.2 × 5.8 μm2 electrodes; 17.8 μm electrode pitch; 20 kHz sampling rate). Acute cerebellar slices were placed flat on the arrays and continuously superfused with carbogen-loaded ACSF (~30-33°C) throughout the experiment. See (Frey et al., 2010) for circuit details of the CMOS-based microelectrode arrays. To identify the locations of the neurons growing over the array, a sequence of overlapping recording configurations was scanned across the array. A configuration is defined as how switches are set to connect a subset of the electrodes to the 126 available readout channels. Switches are re-configurable within a few milliseconds. The largest possible contiguous block includes 6×17 electrodes, and 147 overlapping block configurations were used to scan the whole array at a rate of 1 min per configuration. A recording session lasted 2.5 hours. Recordings were acquired inside an incubator in order to control environmental conditions (36°C and 5% CO2).

### Spatial and temporal upsampling

Signals were up-sampled to 160 kHz using the Nyquist-Shannon sampling theorem and the Whitaker-Shannon interpolation formula to improve temporal resolution (Blanche and Swindale, 2006). The theorem is a mathematical proof that any sampled signal can be perfectly reconstructed into the original analog signal, as long as the analog signal has frequency components less than the Nyquist frequency. The microelectrode array circuitry filters the analog data with a first-order filter with an upper cutoff frequency of 3.7 kHz before data is sampled at 20 kHz, which satisfies the requirements. To estimate waveforms between recording electrodes, up-sampled and averaged EAPs were then spatially interpolated across a 1×1 μm grid by cubic interpolation in Matlab (‘griddata’ function; MathWorks Matlab R2014a).

### Statistics

Pearson correlation and two-side t-tests were done in Matlab R2014a (MathWorks).

## Results

The largest extracellular spike always had negative polarity (Figure 1B) and was typically localized near the proximal AIS or the peak Ankyrin-G (AnkG; a molecular marker of the AIS) immunofluorescence signal (Figure 1A) but never at the soma (Figure 1C). Especially, the signals of cells whose AIS originated from a dendrite (e.g. Figure 1A bottom), clearly evidenced that the EAP was correlated with AIS location and shape. Signals near the soma of such cells typically were not detectable unless averaging multiple EAPs (Figure 1B bottom, electrode 5). An AIS was considered to have a dendritic origin if it began at a neurite branch point, whose other branch expresses microtubule-associated protein 2 (MAP2; a molecular marker of somatodendritic regions) and not AnkG. Out of the 49 neurons from 12 cultures (neurons that may have had overlapping voltage signals were excluded), 26 had an AIS with a dendritic origin, which is a common phenomenon observed in a variety of preparations (Triarhou, 2014). As an ‘axon hillock’ is not obvious for such cells, we define the ‘neck’ as the distance from the proximal end of the AIS to the soma. The neck length was 31 ± 24 μm (mean ± s.d., N = 26) for AISs of dendritic origin and 7 ± 7 μm (N = 23) for AISs of somatic origin. A significant relationship did not exist between the largest EAP spike and neck length (Pearson’s correlation coefficient r = 0.16, p = 0.27, N = 49). Moreover, EAP spike magnitude did not depend on whether the AIS was of somatic or dendritic origin (p = 0.71, two-side t-test, N = 49), which demonstrated that the relative location of the soma does not strongly influence the magnitude of the EAP. Importantly, AIS lengths were positively correlated to spike magnitude (r = 0.28, p = 0.05, N = 49). Soma diameter was positively correlated to the largest EAP spike for AIS with somatic (r = 0.47, p = 0.02, N = 23) but not dendritic (r = 0.06, p = 0.75, N = 26) origins. Overall, AIS lengths and soma diameters were 31 ± 13 μm and 17 ± 5 μm, respectively (mean ± s.d., N = 49).

**Figure 1.**
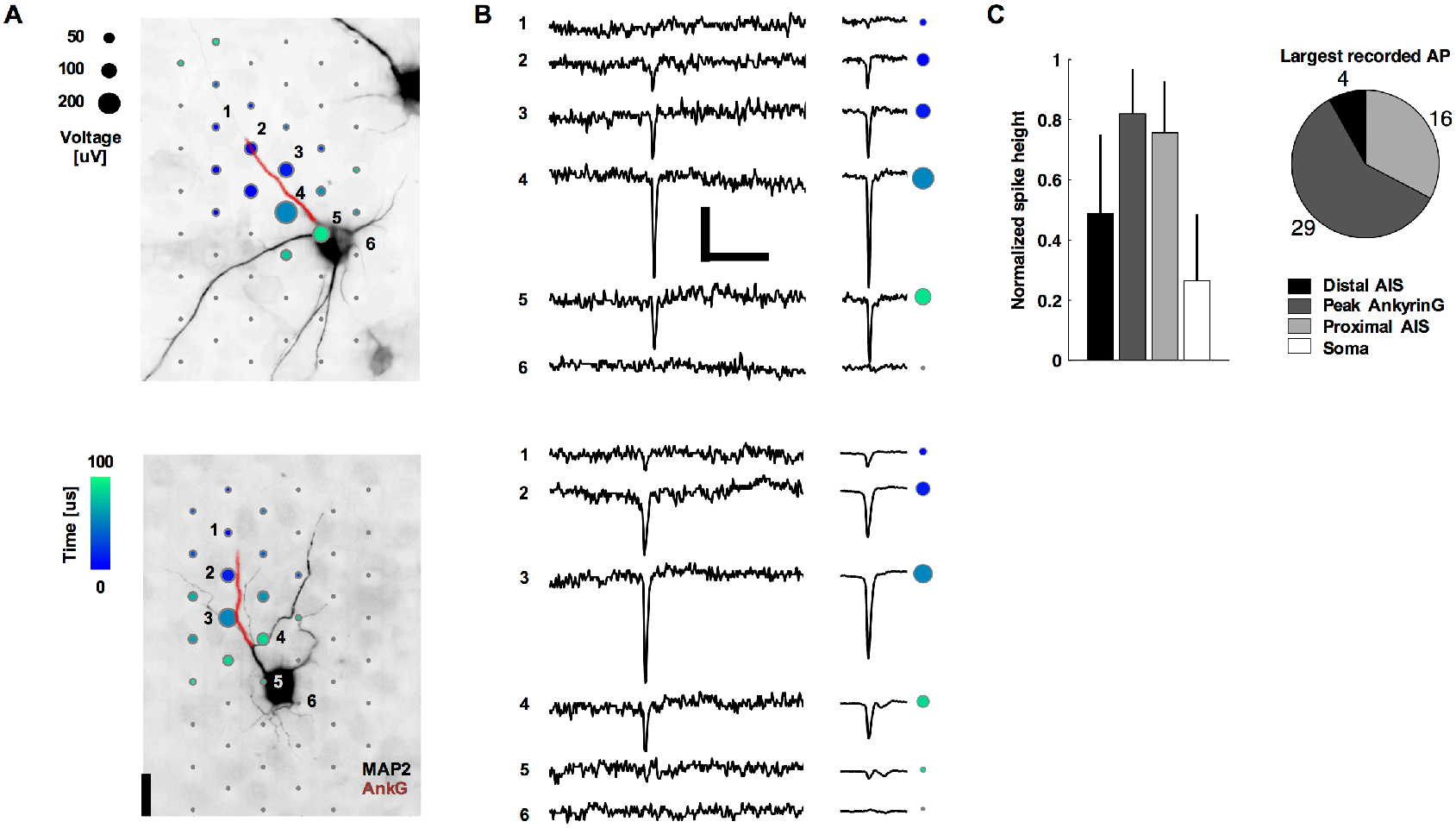
Neurons’ negative-amplitude extracellular action potentials co-localize with their AISs. **A**. Spatial and temporal (color) distribution of the averaged spontaneous extracellular action potential for two neurons. Dots are recording electrode locations, and dot sizes represent the AP negative peak voltage in the EAP. The underlying fluorescence images show somas and dendrites (black; MAP2) and the AIS (red; AnkG). The EAPs are the average of 14 (top neuron) and 200 (bottom) spontaneous spikes. Numbered electrodes correspond to those in B. Scale bar: 20 μm. **B**. Raw voltage waveforms for a single action potential (left) and the averaged waveform as used in A (right; N = 14 top and 200 bottom). Colored dots correspond to those for the same electrodes numbered in A. Scale bars: 100 μV and 5 ms. **C**. Summary statistics of negative spike heights. Data in the bar plots (mean ± s.d., N = 49) are normalized by the peak voltage amplitude in the EAP.

Consistent with most reports in the literature, action potentials typically initiated at the distal AIS (Hu et al., 2009; Baranauskas et al., 2013), and EAP magnitude at the distal AIS was lower than the largest EAP (Figures 1C, 2, 3A-D). Dual-patch recordings of soma and axon (Yu et al., 2008; Baranauskas et al., 2013), voltage sensitive dye recordings (Palmer and Stuart, 2006), and modeling (Kole et al., 2008; Baranauskas et al., 2013) show that action potentials initiate in the axon at about 40 μm from the soma. This is consistent with the location of the distal end of the AIS we found for AISs originating from the soma (36 ± 13 μm, mean ± s.d.). Once initiated, action potentials propagated in both directions: into the axon proper and backwards through the proximal AIS and towards the soma. This scenario holds for both cases of axons originating directly from the soma or from dendrites (Figure 2). Of the 22 neurons with an AIS length longer than 30 μm, the propagation velocity was 1.1 ± 1.5 m/s (mean ± s.d.; 65.6 [im and 0.6 m/s median); the distal end of the AIS always fired first. The velocities were similar to those of action potentials propagating in unmyelinated cortical axons (Bakkum et al., 2013). Some of the neurons with shorter AIS lengths showed action potentials occurring nearly simultaneously throughout the AIS or, in 2 cases, first at the proximal end of the AIS or at the soma (Figure 3D). For the 15 neurons with a neck length greater than 30 μm, the propagation velocity from the proximal AIS to the soma center was slower at 0.3 ± 0.2 m/s (mean ± s.d.; 0.2 m/s median).

**Figure 2.**
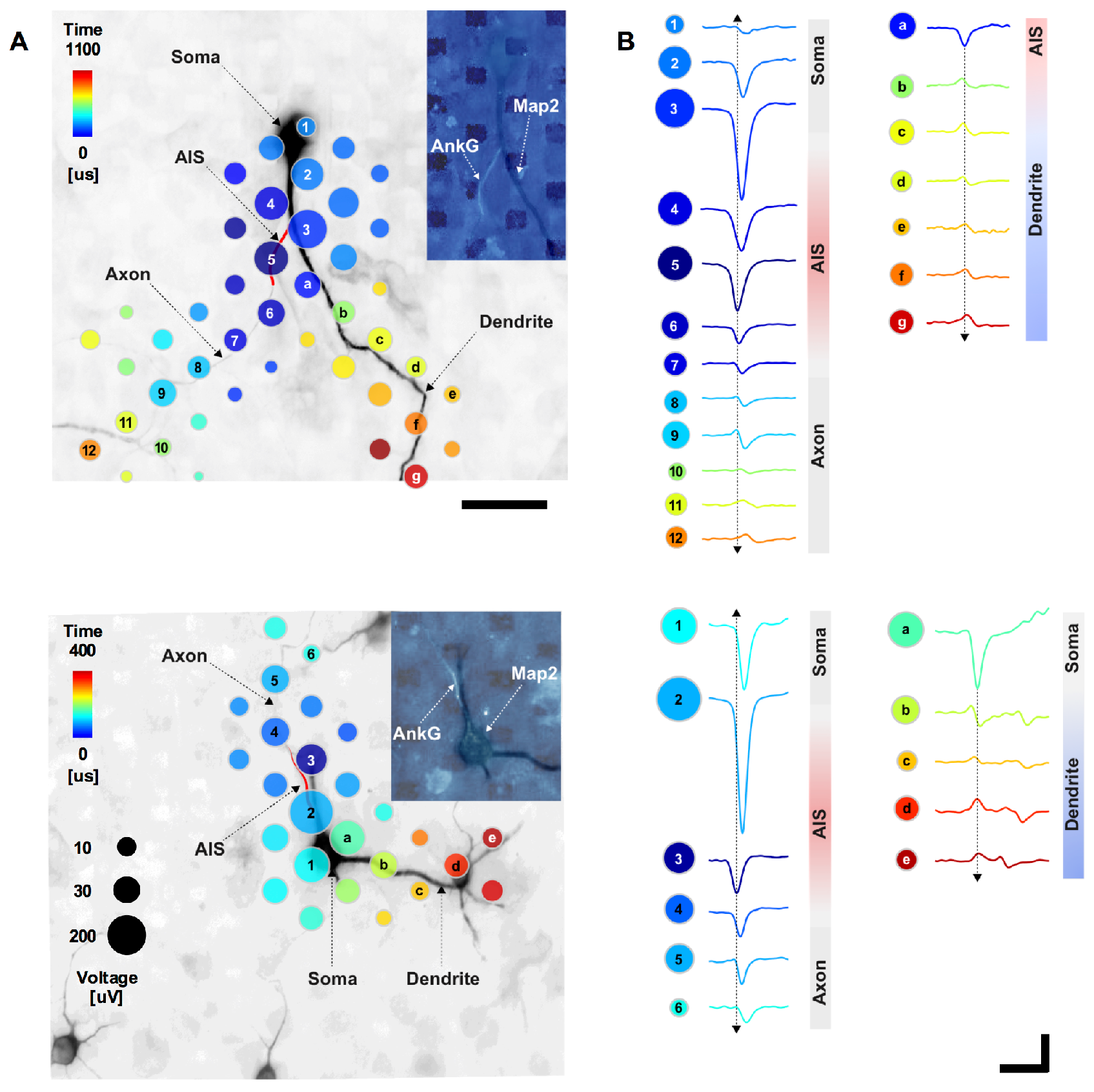
Detailed EAP. **A**. Spatial and temporal distribution of the averaged spontaneous EAP for a neuron with an AIS originating from a dendrite (top) and an AIS originating from a soma (bottom). Dots mark recording-electrode locations. Time (dot color) is the delay from the first voltage peak, detected at the distal AIS, until the peak absolute voltage signal (dot size) at each location. Axonal and somatic areas correspond to signals that had larger negative peaks, while the dendritic area corresponds to signals that had larger positive peaks. The underlying fluorescence image shows somas, dendrites (black; MAP2), and the AIS (red cartoon; AnkG). The inset fluorescence image shows MAP2 (dark) and AnkG (light); dark rectangles are electrodes. Scale bar: 50 μm. **B**. Average voltage waveforms recorded at the electrodes with matching labels to those in A (i.e. numbers and letters). Arrows indicated signal propagation direction. Scale bars: 100 μV vertical; 1 ms horizontal.

As a control, live-cell imaging and patch clamp electrophysiology verified that the largest-amplitude extracellular action potential is spatially offset from the soma and distally initiated (Figure 3E-F). Here, action potentials were evoked by applying current injection through the patch pipette. Results were consistent across five experiments. Alexa Fluor 594 dye was injected into the cell via the patch pipette in order to perform simultaneous live-cell imaging. The extracellular signal is approximately proportional to the negative derivative of the intracellular signal (Henze et al., 2000) (and is proportional to the membrane current) (Henze et al., 2000; Gold et al., 2006; Einevoll et al., 2013). Then the peak negative extracellular amplitude corresponds to the point of maximum positive slope on the intracellular trace (Figure 3F). The EAP is ~2 orders of magnitude smaller than an intracellular AP. Therefore, periods of slower change in the intracellular AP correspond to a small or not detectable EAP. The initial positive deflection in the EAP near the soma is typical of capacitive membrane currents. These would be caused by intracellular currents, spreading axially from the AIS, charging the membrane prior to the opening of voltage gated ion channels.

**Figure 3.**
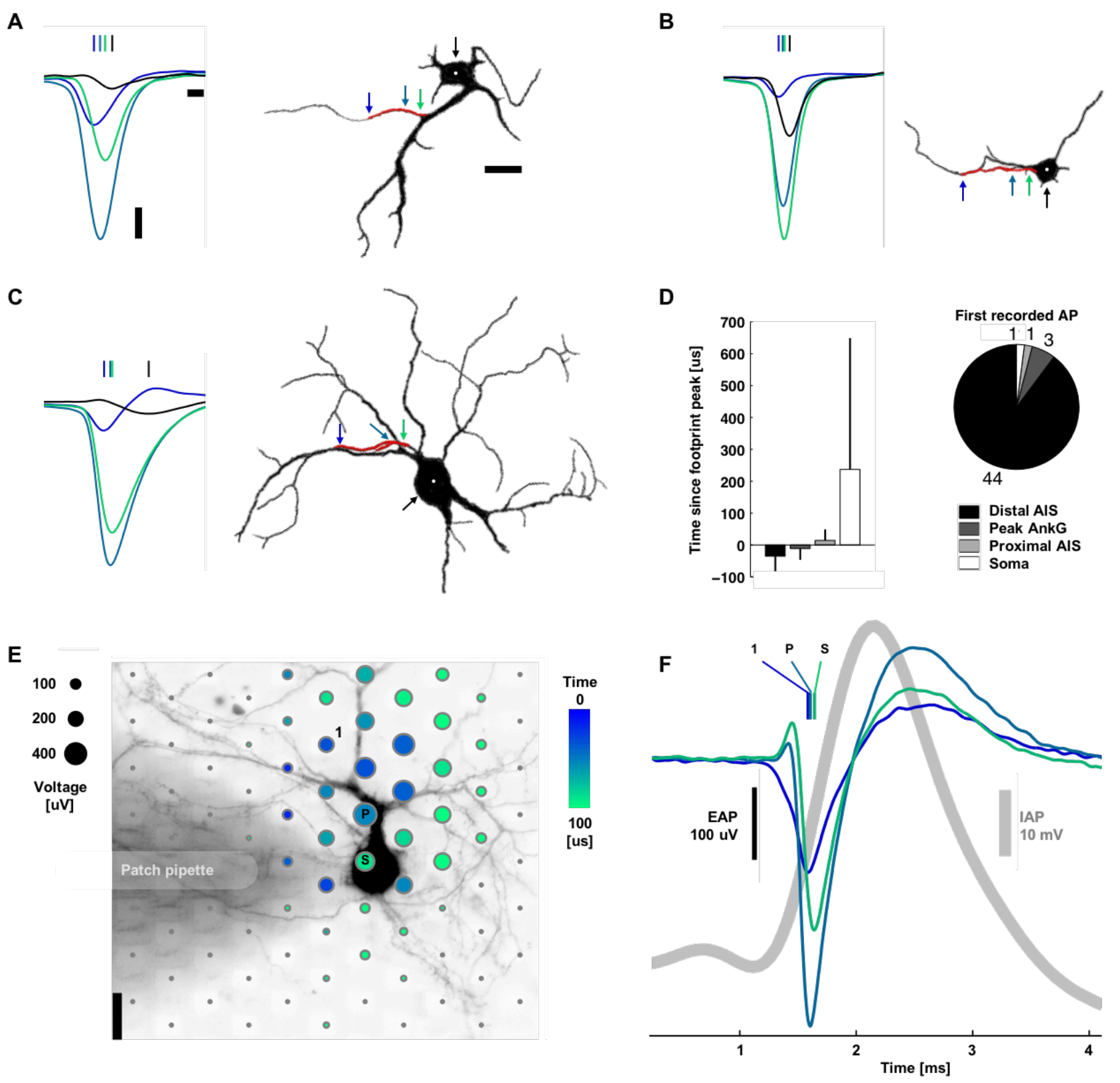
Action potentials typically initiated at the distal AIS. **A**, **B**, **C**. Left: Spontaneous action potentials at the distal AIS (blue), peak ankyrin-G fluorescence signal (blue-green), proximal AIS (green), and soma (black). Vertical lines indicate the timing of the negative voltage peak voltage. Waveforms are the average of 11, 14, and 69 spikes for A, B, and C. Right: Silhouettes of immunostained neurons’ fluorescence images (black: MAP2, red: ankyrin-G). Colored arrows and the white dot indicate the location of the plotted waveforms. Scale bars: 100 μV vertical; 100 μs left; 20 μm right. **D**. Summary statistics. Bar plots are mean ± s.t.d, N = 49. **E**. Spatial and temporal (color) distribution of the averaged EAP (dot sizes). The shaded area is a patch pipette that evoked action potentials by current injection and delivered the fluorescent dye for live-cell imaging. The labeled electrodes correspond to those in B: the 1st electrode recording an action potential, the electrode recording the Peak voltage signal, and the electrode nearest the Soma. Scale bar: 20 μm. **F**. Average intracellular (gray) and extracellular (color) action potential waveforms (39 spikes). Colored vertical lines indicate the timing of the negative voltage peak.

While the largest amplitude spikes had negative polarity and were colocalized with AISs, spikes with positive polarity were co-localized with some, but not all, dendritic branches (Figure 4A). The average peak negative spike was 6.9 times bigger than the average peak positive spike and 5.9 times bigger than the average somatic spike, which also had a negative polarity (N = 24) (Figure 4C). The co-localization of the positive peaks with dendrites and their mostly monophasic nature suggest that they are caused by a passive (i.e., capacitive) return current balancing the influx of sodium ions at the AIS. Alternatively, voltage-gated potassium channels are expressed in dendrites and could also be involved (Korngreen and Sakmann, 2000; Martina et al., 2003). Interestingly, in our measurements (Figure 4B) we found a delay between the occurrence of negative and positive spikes: 113 ± 108 μs (mean ± s.d., 86 μs median, N = 24; see also Movie 1). The delay was significantly correlated to axial distance (r = 0.79, p = 0.001; Figure 4B) and not straight-line distance (r = 0.34, p = 0.11), suggesting an intracellular influence (Destexhe and Bedard, 2012). An ideal return current would result in simultaneous peaks. However, signals at these locations will be influenced by other components, such as active ionic currents from nearby neurites and the presence of other cells, such as glia, that may affect resistivity. Likewise, how and why dendritic branches differentially contribute to the EAP (Figure 4A) is an important future study; possibly, the intracellular resistance varies for different dendritic branches. For Figure 4, 25 of the 49 neurons were excluded because either some distal parts of their dendritic arbor were not recorded at the same time as the AIS or their dendritic arbor overlapped with neurites from another cell.

**Figure 4.**
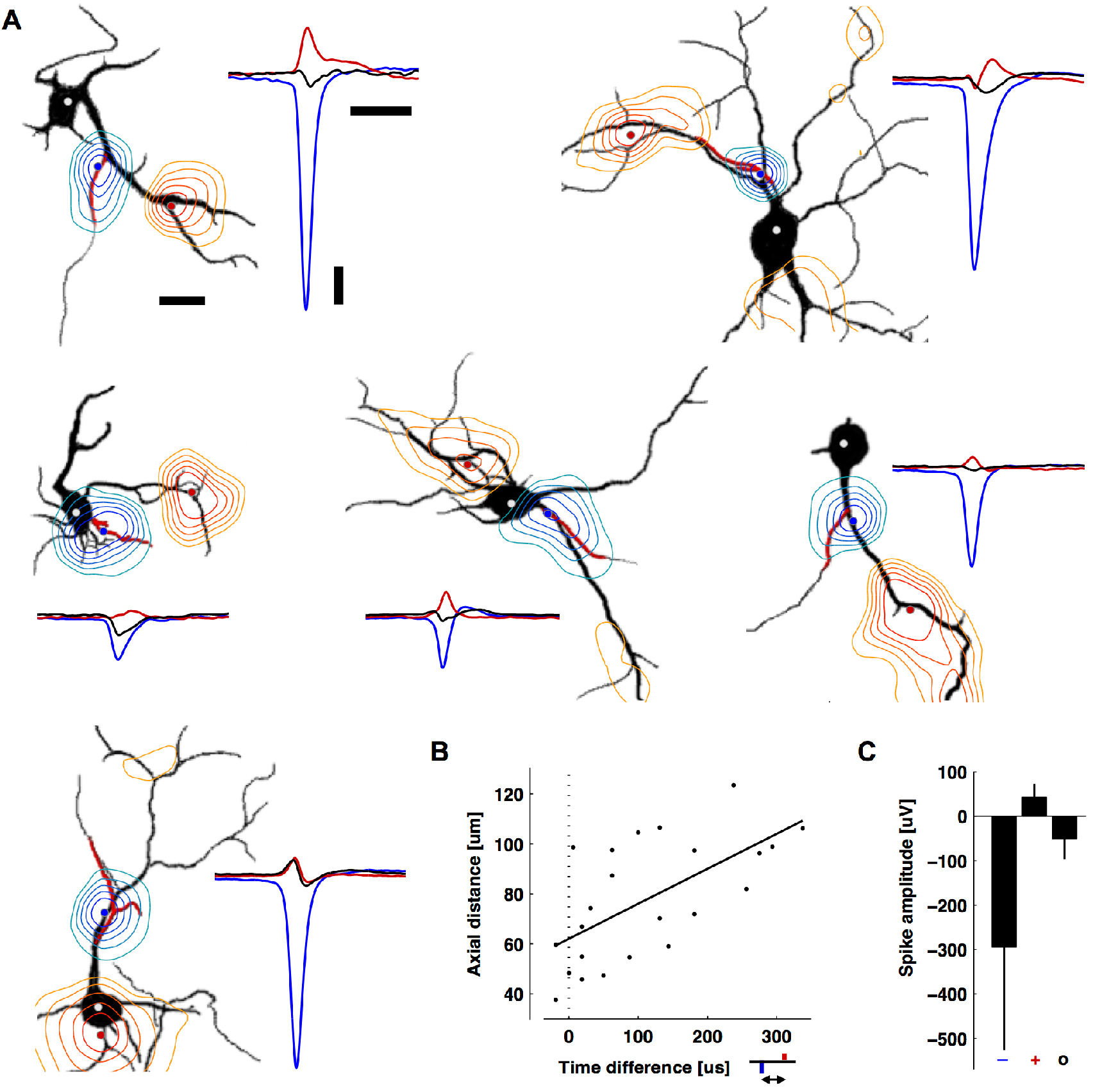
Positive-amplitude extracellular action potentials co-localized with some, but not all, dendritic branches and occurred after the AIS spike. **A**. Contour plots of the minimum (blue) or maximum (red) EAP within ± 500 μs of the negative peak (black silhouettes: MAP2, red: ankyrin-G). The contours are normalized to the largest negative signal (blue-to-green) or the largest positive signal (red-to-yellow). The blue, red, and white (soma) dots are the locations of the plotted waveforms. The two neurons in the top row are also in Figure 3. From left to right and top to bottom, EAPs are averages of 11, 69, 28, 68, 52, and 199 spontaneous spikes. Scale bars: 100 μV vertical; 1 ms right; 20 μm left. **B**. The time delay versus axial distance between the largest negative peak (blue) and the largest positive peak (red) voltages in the EAP. The line is a linear regression (Pearson’s correlation coefficient r = 0.79, p = 0.001, N = 24). **C**. Average peak negative, peak positive, and somatic spike amplitudes (mean ± s.d., N = 24). Time courses of the EAPs are seen in Movie 1.

To investigate the role of the AIS also in neuronal tissue closely resembling in-vivo morphology, we performed further experiments in Purkinje cells in acute cerebellar mouse slices. Purkinje cells differ from cortical neurons by having a more passive dendritic tree and larger soma. These two features would make it more likely that the largest extracellular signal would arise from the soma, if the soma instead of the AIS would be the source of the signal. Nevertheless, corroborating the results from cortical cell cultures above, the largest negative EAPs (Figure 5A) occurred at the area containing axons instead of at the Purkinje cell layer (i.e., the location of all Purkinje somas), and the positive EAPs (Figure 5B) consistently occurred in the area containing dendrites. In all cases, the largest negative EAPs were found to co-localize with the putative location of the Purkinje cell AISs; the mean of negative amplitudes peaked sharply approximately 40 μm away from the center of the Purkinje cell layer, in the granular cell layer (Figure 5C). On the other hand, the largest positive EAPs (i.e., the return currents) occurred 40 to 100 μm away from the center of the Purkinje cell layer, in the molecular layer. Purkinje cells have a stereotypical orientation: their AISs are always at the granular cell layer, between the white matter and the Purkinje cell layer; their dendritic arbors are located in the molecular layer, fanning out opposite the AIS from the Purkinje cell layer towards the pia. Parasagittal slices were made in order to keep the full morphology of Purkinje cells intact for recording. Purkinje cells are GABAergic and produce large spontaneous spikes even without synaptic input; the parallel fibers that would provide input get cut during slicing.

**Figure 5.**
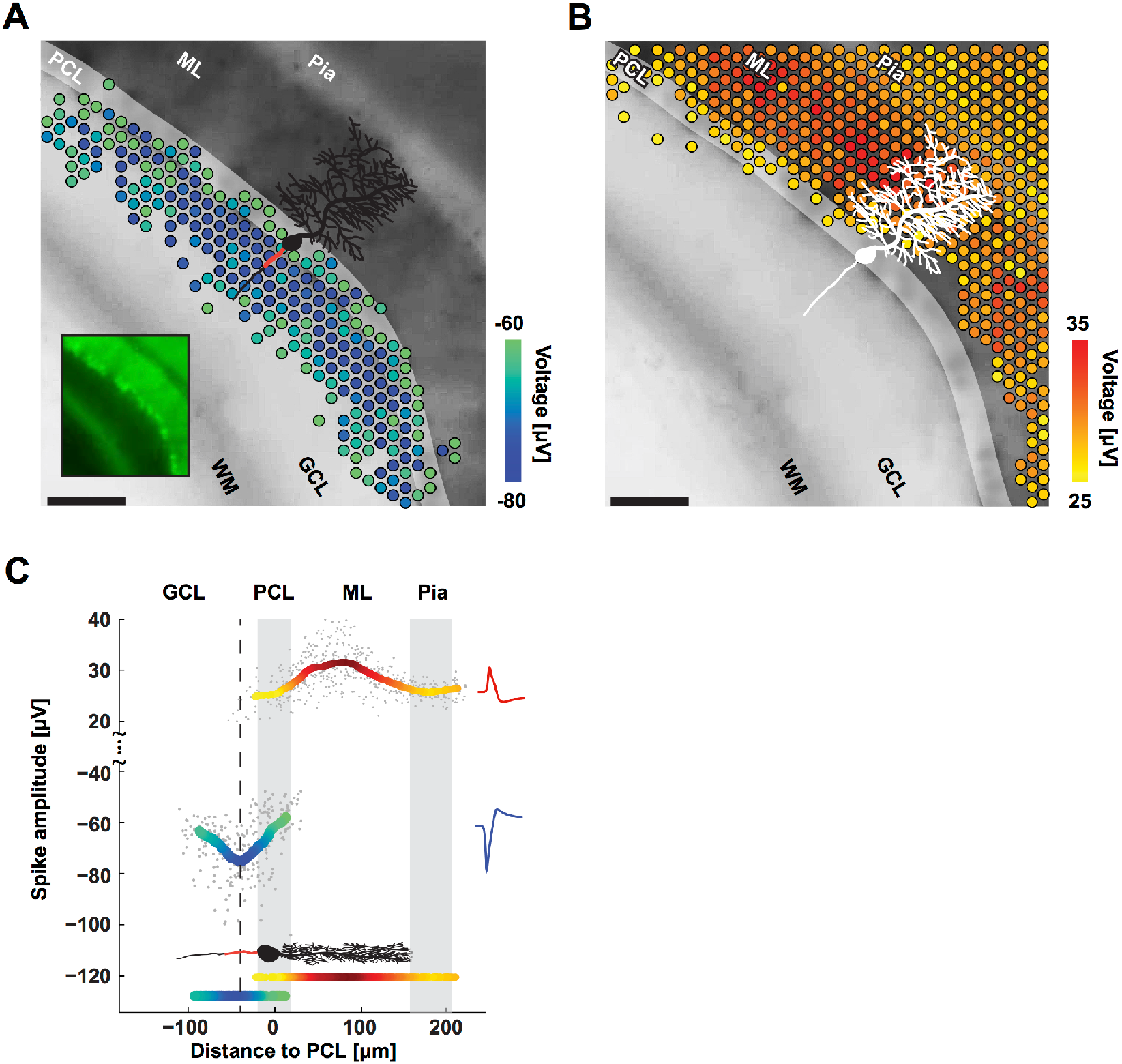
Purkinje cells’ negative and positive EAPs co-localize with AISs and dendrites. **A**. Spatial distribution of the most negative EAPs recorded from an acute cerebellar slice. Dots show the locations of recording electrodes that detect EAPs (color) below a −60 μV threshold. EAP amplitudes are the mean voltage of the 20 largest events during a 10 second recording. The underlying black-and-white live-cell fluorescence image, taken immediately after recording, shows the location of the cerebellar layers: white matter (WM), granular cell layer (GCL), Purkinje cell layer (PCL), molecular layer (ML), and pia. The somas of the Purkinje cells are in the PCL (highlighted for clarity; easier to see in the smaller inset image). All Purkinje cell dendrites are in the ML, all AISs are in the GCL. The microscopy image captured cells at the top layer of the acute slice, while the EAPs were recorded from cells at the bottom layer. A sample cartoon of a Purkinje cell is shown in black with a red line estimating the AIS location. Scale bar: 100 μm. **B**. Same as A except for EAPs above a 25 μV threshold. **C**. Summary of EAP amplitudes. Gray dots represent the largest negative or positive EAP amplitudes detected at each electrode versus the distance of the electrode from the PCL. Colored curves represent the mean for both negative (blue-green) and positive (yellow-red) EAPs as measured by the respective electrodes. The minimum in the curve representing the negative EAPs (dashed line at −40 μm) co-localizes with the putative location of AISs (red line in the cartoon of a Purkinje cell) in the GCL. The largest positive EAPs co-localize with the dendrites in the ML. Representative spike shapes of the negative (blue) and positive (red) EAPs recorded from the acute cerebellar slice are shown on the right. Movie 1. The videos give contour plots of the EAP from neurons in Figure 4. While voltages in Figure 4 are the peak values within a ±500 μs time range, voltages here are measured at the time indicated in the movie text. The background is a MAP2 fluorescence image with a drawn red line to indicate the AIS locations. Blue contours are negative voltages, and red contours are positive voltages. Adjacent contour lines are separated by −50 μV (blue; starting from −10 μV) or +10 μV (red; starting from +10 μV). Scale bar: 20 μm.

## Discussion

Extracellular signal amplitudes can be influenced by many factors, such as electrode impedance, the local microenvironment, or the degree of coupling between a neuronal membrane and an electrode. However, these influences likely had a negligible effect on our results. We deposited platinum black on the electrodes (see Methods). Besides reducing the electrode impedance, platinum black also reduces variations in signal amplitude due to impedance mismatches across the arrays’ electrodes to about 5% (Viswam et al., 2014). This variation is negligible when compared to the AIS-to-soma or AIS-to-dendrites signal amplitudes that differ by over 600% on average (Fig. 4C). Consistent results across 49 individual neurons and multiple preparations provide strong evidence that the results were not an artifact from variations in local microenvironments or coupling. Further evidence is that a neuron’s EAP is detected by multiple neighboring electrodes, including sites that are not immediately adjacent or coupled to a neuronal membrane, and that the signal attenuates with increasing distance from the source as expected (Figs. 1A, 2A, 3E, 4A).

The general phenomenon that the largest signal amplitudes occur in the region of the AIS is applicable to both in vitro and in vivo situations. The EAP arises from neuronal membrane currents, in particular from voltage-gated sodium channels. Previous studies showed that the molecular composition of axons in vitro and in vivo, including sodium channel distribution in the AIS, are consistent (Hedstrom et al., 2007). This implies that the membrane currents responsible for the EAP will likewise be similar. In turn, how the electric field spreads in the medium and is picked up as a potential on the recording electrodes is governed by laws of physics, i.e., Maxwell’s equations (Einevoll et al., 2013). In other words, this aspect is independent of whether the experimental preparation is in vitro or in vivo. Nevertheless, we also experimentally addressed this issue and found consistent results in acute cerebellar slices, which preserve in vivo local microarchitecture (Fig. 5).

The basic and fundamental observation of the EAP being driven by currents from the AIS instead of the soma can affect how extracellular recordings are interpreted. For example, neurons with spherically symmetric dendrites are thought to produce small EAPs due to the summation of dendritic membrane currents canceling each other out (axonal contributions presumed negligible) (Logothetis, 2008; Buzsaki et al., 2012). However, such symmetry is broken when membrane currents originate from the AIS or from specific dendritic branches (Figure 4A). Likewise, neurons with smaller soma are thought to produce smaller EAPs (Humphrey and Schmidt, 1990), but we observed this only for neurons with an AIS originating from the soma. Why wide variations in EAP waveforms exist between cells has been unclear (Anastassiou et al., 2015) but can be explained to a degree by variations in AIS shape and level of AIS formation. In our preparation, EAP magnitudes ranged between 67 to 952 μV (up to 1.8 mV in previous data (Bakkum et al., 2013)) and AIS lengths between 13 to 71 μm. ‘Silent’ cells existed and were typically associated with decreased or no AnkG expression. Occasionally, AISs were forked, which produced distorted or double-spiked EAPs depending on the recording location. Why different dendritic branches produce different extracellular signals (cf. Figure 4A) is harder to explain and may require different experimental approaches to decipher. In tangential work, the AIS has been identified as the site with the lowest threshold and highest reliability for extracellular stimulation (McIntyre and Grill, 1999; Fried et al., 2009; Radivojevic et al., 2016), whereas stimulation under the soma often failed to evoke action potentials (Radivojevic et al., 2016). This finding supports the AIS’s higher contribution to the EAP and provides further indirect evidence that the AIS contains a higher concentration of voltage-gated ion channels. In summary, the action’s at the AIS.

## Additional information

### Author Contributions

Experimental design and data analysis was done by D.J.B., D.J., and M.E.J.O.; D.J.B., D.J., M.E.J.O., and M.R. performed experiments; technical support was provided by D.J.B., D.J., U.F., and M.R.; D.J.B., U.F., M.E.J.O., A.H., and H.T. wrote the manuscript; and project supervision was done by U.F., H.T. and A.H. All authors approve the final version of the manuscript and agree to be accountable for all aspects of the work in ensuring that questions related to the accuracy or integrity of any part of the work are appropriately investigated and resolved. All listed authors qualify for authorship, and all those who qualify for authorship are listed.

## Acknowledgments

We thank Alexander Stettler and Albert Martel for postprocessing CMOS chips; Felix Franke and Wei Gong for critical discussions; Jan Muller for technical support; and the D-BSSE staff for expediting experiments.

## References

Anastassiou CA, Perin R, Buzsaki G, Markram H, Koch C (2015) Cell type- and activity-dependent extracellular correlates of intracellular spiking. J Neurophysiol 114:608–623.

Bakkum DJ, Frey U, Radivojevic M, Russell TL, Muller J, Fiscella M, Takahashi H, Hierlemann A (2013) Tracking axonal action potential propagation on a high-density microelectrode array across hundreds of sites. Nat Commun 4:2181.

Baranauskas G, David Y, Fleidervish IA (2013) Spatial mismatch between the Na+ flux and spike initiation in axon initial segment. Proc Natl Acad Sci U S A 110:4051–4056.

Blanche TJ, Swindale NV (2006) Nyquist interpolation improves neuron yield in multiunit recordings. J Neurosci Methods 155:81–91.

Buzsaki G, Anastassiou CA, Koch C (2012) The origin of extracellular fields and currents-EEG, ECoG, LFP and spikes. Nature Reviews Neuroscience 13:407–420.

Colbert CM, Johnston D (1996) Axonal action-potential initiation and Na+ channel densities in the soma and axon initial segment of subicular pyramidal neurons. J Neurosci 16:6676–6686.

Destexhe A, Bedard C (2012) Do neurons generate monopolar current sources? J Neurophysiol 108:953–955.

Dodge FA, Cooley JW (1973) Action potential of the motorneuron. IBM Journal of Research and Development 17:219–229.

Einevoll GT, Kayser C, Logothetis NK, Panzeri S (2013) Modelling and analysis of local field potentials for studying the function of cortical circuits. Nat Rev Neurosci 14:770–785.

Frey U, Egert U, Heer F, Hafizovic S, Hierlemann A (2009) Microelectronic System for High-Resolution Mapping of Extracellular Electric Fields Applied to Brain Slices. Biosensors and Bioelectronics 24:2191–2198.

Frey U, Sedivy J, Heer F, Pedron R, Ballini M, Mueller J, Bakkum D, Hafizovic S, Faraci FD, Greve F, Kirstein KU, Hierlemann A (2010) Switch-Matrix-Based High-Density Microelectrode Array in CMOS Technology. Ieee Journal of Solid-State Circuits 45:467–482.

Fried SI, Lasker AC, Desai NJ, Eddington DK, Rizzo JF, 3rd (2009) Axonal sodium-channel bands shape the response to electric stimulation in retinal ganglion cells. J Neurophysiol 101:1972–1987.

Gold C, Henze DA, Koch C, Buzsaki G (2006) On the Origin of the Extracellular Action Potential Waveform: A Modeling Study. J Neurophysiol 95:3113–3128.

Grace AA, Bunney BS (1983) Intracellular and extracellular electrophysiology of nigral dopaminergic neurons--2. Action potential generating mechanisms and morphological correlates. Neuroscience 10:317–331.

Hallermann S, de Kock CP, Stuart GJ, Kole MH (2012) State and location dependence of action potential metabolic cost in cortical pyramidal neurons. Nat Neurosci 15:1007–1014.

Hedstrom KL, Xu X, Ogawa Y, Frischknecht R, Seidenbecher CI, Shrager P, Rasband MN (2007) Neurofascin assembles a specialized extracellular matrix at the axon initial segment. J Cell Biol 178:875–886.

Henze DA, Borhegyi Z, Csicsvari J, Mamiya A, Harris KD, Buzsaki G (2000) Intracellular Features Predicted by Extracellular Recordings in the Hippocampus In Vivo. J Neurophysiol 84:390–400.

Holt GR, Koch C (1999) Electrical interactions via the extracellular potential near cell bodies. Journal of computational neuroscience 6:169–184.

Hu W, Tian C, Li T, Yang M, Hou H, Shu Y (2009) Distinct contributions of Na(v)1.6 and Na(v)1.2 in action potential initiation and backpropagation. Nat Neurosci 12:996–1002.

Humphrey DR, Schmidt EM (1990) Extracellular single-unit recording methods. In: Neurophysiological techniques, pp 1–64: Humana Press.

Kole MH, Ilschner SU, Kampa BM, Williams SR, Ruben PC, Stuart GJ (2008) Action potential generation requires a high sodium channel density in the axon initial segment. Nat Neurosci 11:178–186.

Korngreen A, Sakmann B (2000) Voltage-gated K+ channels in layer 5 neocortical pyramidal neurones from young rats: subtypes and gradients. J Physiol 525 Pt 3:621–639.

Logothetis NK (2008) What we can do and what we cannot do with fMRI. Nature 453:869–878.

Lorincz A, Nusser Z (2010) Molecular identity of dendritic voltage-gated sodium channels. Science 328:906–909.

Mainen ZF, Joerges J, Huguenard JR, Sejnowski TJ (1995) A model of spike initiation in neocortical pyramidal neurons. Neuron 15:1427–1439.

Martina M, Yao GL, Bean BP (2003) Properties and functional role of voltage-dependent potassium channels in dendrites of rat cerebellar Purkinje neurons. J Neurosci 23:5698–5707.

McIntyre CC, Grill WM (1999) Excitation of central nervous system neurons by nonuniform electric fields. Biophysical Journal 76:878–888.

Palmer LM, Stuart GJ (2006) Site of action potential initiation in layer 5 pyramidal neurons. J Neurosci 26:1854–1863.

Petersen AV, Johansen EO, Perrier JF (2015) Fast and reliable identification of axons, axon initial segments and dendrites with local field potential recording. Front Cell Neurosci 9:429.

Pettersen KH, Einevoll GT (2008) Amplitude variability and extracellular low-pass filtering of neuronal spikes. Biophysical Journal 94:784–802.

Radivojevic M, Jackel D, Altermatt M, Muller J, Viswam V, Hierlemann A, Bakkum DJ (2016) Electrical Identification and Selective Microstimulation of Neuronal Compartments Based on Features of Extracellular Action Potentials. Sci Rep 6:31332.

Rall W (1962) Electrophysiology of a dendritic neuron model. Biophysical Journal 2:145–167.

Schmidt-Hieber C, Bischofberger J (2010) Fast sodium channel gating supports localized and efficient axonal action potential initiation. J Neurosci 30:10233–10242.

Tamamaki N, Yanagawa Y, Tomioka R, Miyazaki J, Obata K, Kaneko T (2003) Green fluorescent protein expression and colocalization with calretinin, parvalbumin, and somatostatin in the GAD67-GFP knock-in mouse. J Comp Neurol 467:60–79.

Triarhou LC (2014) Axons emanating from dendrites: phylogenetic repercussions with Cajalian hues. Front Neuroanat 8:133.

Viswam V, Jackel D, Jones I, Ballini M, Muller J, Stettler A, Frey U, Franke F, Hierlemann A (2014) Effects of Sub-10μm Electrode Sizes on Extracellular Recording of Neuronal Cells. In: 18th International Conference on Miniaturized Systems for Chemistry and Life Sciences (MicroTAS 2014), pp 980–982. San Antonio, Texas.

Yu Y, Shu Y, McCormick DA (2008) Cortical action potential backpropagation explains spike threshold variability and rapid-onset kinetics. J Neurosci 28:7260–7272.

